# The choreography of human attraction: physiological synchrony in a blind date setting

**DOI:** 10.1101/748707

**Authors:** E. Prochazkova, E. E. Sjak-Shie, F. Behrens, D. Lindh, M. E. Kret

## Abstract

Humans are social animals whose mental wellbeing is shaped by the ability to attract and connect with each other. In a dating world, in which success can be determined by brief interactions, apart from physical features, there is a whole choreography of movements, physical reactions and subtle expressions that drive humans’ sexual attraction. To determine what drives attraction, we measured nonverbal dynamics between people during real-life interactions outside the laboratory, where dating is most relevant. Participants wore eye-tracking glasses with embedded cameras, and devices to measure physiological signals including heart rate and skin conductance. Crucially, visible signals that can be controlled, such as facial expressions or gaze, did not predict attraction. Instead, attraction was predicted by synchrony in heart rate and skin conductance between partners, which are unconscious and difficult to regulate. Our findings suggest that shared emotionality is vital for mutual attraction. Moreover, physiological synchrony may provide a medium for translating visible expressions into embodied emotions, which can turn into intentions via somatosensory simulation.

**Significance Statement:** In our modern world where millions of people meet online without interacting face-to-face, the question “what defines attraction” has become very relevant. In this study, we used modern technologies to lay down a foundation for the processes that drive human attraction during real-life interactions. Contrary to common belief, we found that attraction is not predicted by a frequency of expression or eye fixation duration, nor is linearly related to the participant’s autonomic nervous system activity. Importantly, we found that the more a participant synchronized their heart rate and skin conductance with their partner, the more they felt attracted toward that person. This study reveals a fundamental physiological process that plays a role in the formation of romantic relations.

## Introduction

In the past decade, we have witnessed rapid changes in the modern dating culture. With more than 50 million people dating online, of whom 80% are looking for a serious relationship, and more than 1.5 million Tinder dates per week (1), online dating has become a cultural force that shapes the way our generation interacts with and relates to each other (2). Consequently, three main vicissitudes have occurred: First, in contrast to previous generations, dates happen largely between strangers. Second, dating decisions are made upon short interactions. Third, the access to many potential partners increases the difficulty of choosing one partner. In the social realm, this global phenomenon can be placed in the context of an even broader fundamental puzzle: In the world of vast dating possibilities, what drives feelings of attraction between people? We took a data-driven approach to answer this question. In this blind date experiment, we used state of the art technology including eye-tracking glasses linked to physiological measures to track a whole choreography of movements, subtle expressions and physiological reactions to predict sexual attraction between people.

Physical attractiveness is often valued as one of the most important characteristics of a potential partner (3). Yet, research demonstrates that judging a potential romantic partner based on a photo does not predict how attractive this person will be rated after a social interaction (4). In a social situation, apart from verbal information, a variety of nonverbal dynamics such as eye gaze, facial expressions and gesticulations influence attraction. For instance, both smiling and laughing have been reported to reflect the degree of attraction to the other person, and subsequently cause the other to be attracted to the person expressing the smiles and laughter (5–7). Similarly, head nods and open body postures are associated with more self-reported feelings of love among newly met and long-term committed relationship partners (8–10). In addition, dominant body posture, direct eye contact (5, 11, 12), increased skin conductance and heart rate responses have all been linked to the perception of more attractive, opposite-sex targets (13–15). People are often unaware of being influenced by others’ affective displays. This is evident from studies showing that friends and lovers implicitly mimic each other’s nonverbal behavior, such as gaze and facial expressions (16–18). Apart from visible expressions, a series of recent experiments have demonstrated that committed romantic partners synchronize their heart rate and skin conductance. Crucially, the level of synchrony was positively associated with the amount of time spent together and the quality of the relationship (19–21). This suggests that apart from nonverbal signals, physiological synchrony shapes social perception and promotes pro-social behavior.

Taken together, the current literature suggests that attraction emerges from the dynamic exchange of verbal and nonverbal signals (5, 9–11, 22), yet the necessary empirical and analytic tools to directly address this hypothesis were not available until recent times. Consequently, the direct link between nonverbal behavior, physiology and attraction has never been directly verified. To define what drives feeling of attraction, we built a dating lab outside of the regular laboratory setting, where meeting a new person is most natural (Fig. 1). Males and females, who had never met before, entered the dating cabin and had a short date consisting of verbal and nonverbal interactions. We put forward two main hypotheses: The first hypothesis was based on the assumption that individuals’ nonverbal signals and physiology reflect honest affective intentions (23, 24). For example, visible expressions such as frequent smiling, laughing, head nodding and hair touching may give away that a person feels attracted to someone. Furthermore, given the strong link between the physiology and emotions (11, 25, 26), it should be possible to predict attraction purely by looking at participants’ behavioral and physiological responses. In contrast, the second hypothesis was based on the theory that emotional expressions do not communicate emotions - they moderate social exchange (27–29). If true, in order to predict attraction, we cannot focus on individuals’ expressions alone; instead we need to focus on whether expressions are synchronously exchanged between individuals. In the second approach, we took the analysis one step further and measured how synchrony between couples’ eye gaze, expression and physiology influences changes in participants’ attraction.

**Figure 1.**
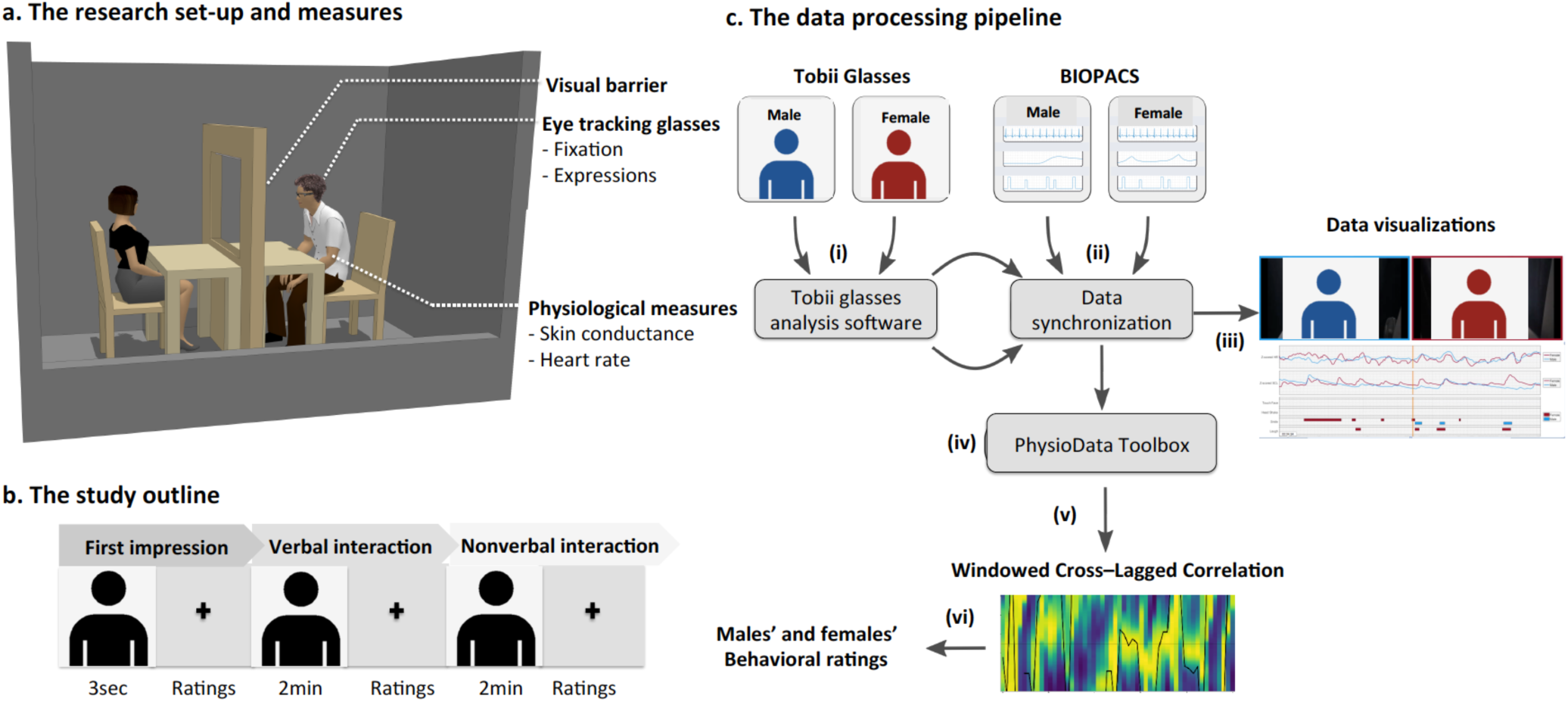
**(a) The experimental set-up** was situated in a habitable container. Inside the cabin, there was a table with two chairs on opposite sides. A white barrier with a fixation cross was placed in the middle of the table, preventing the dyad from seeing each other and controlling the dating interaction. Participants were instructed to remain silent until they heard pre-recorded instructions via a speaker. Throughout the experiment, Tobii eye-tracking glasses measured subjects’ gaze fixations and expressions while participants’ physiology was recorded with two BIOPACs. **(b) Experimental outline.** Males and females, who had never met before, entered the dating cabin and sat at a table. A visual barrier occluded their view, but then opened for three seconds, allowing partners’ to form a first impression. This was followed by one verbal and one nonverbal interaction of 2 minutes each, the order of which was counterbalanced (interactions preceded by a 30 seconds closed barrier baseline). After each interaction, the barrier closed and subjects rated their partner on attraction (0 – 9 point scale). At the end of the experiment, participants could decide whether they wanted to go on another date with their partner and indicate whether they thought their partner wanted to date them. **(c) Pre-processing pipeline.** (i) Two groups of independent coders rated behavioral expressions, and mapped eye gaze fixations on pre-selected areas of interest (ii) gaze fixations and expressions were time locked and synchronized with physiological measures (heart rate, skin conductance) using customized scripts (iii) video visualizations were created (iv) the physiological data were further pre-processed with our PhysioData Toolbox (30) and down-sampled to 100 ms windows for further (v) Windowed Cross-Lagged Correlation analyses (31) before they were (vi) regressed with attraction ratings.

## Results

A dating experiment provides an excellent scenario to test how people infer others’ internal states. This is because during dating interactions, people are likely to exchange a broad variety of facial expressions and gestures, which can help to make inferences about a partner’s romantic intentions. Before we analyzed specific behavioral or physiological pattern that predicts attraction, we examined how accurate people are at predicting whether or not a dating partner wants to date them again in the future.

### How accurate are people at predicting their partner’s romantic interest?

To test participants’ ability to predict their partner’s romantic interest, we asked subjects at the end of the experiment whether they thought their partner would want to date them again (yes/no). Surprisingly, only about half of the subjects (54%) correctly predicted their partner’s answers, which means participants’ accuracy was at chance level (*χ*^*2*^(1) = 1.06, *p* = 0.30, females: 56%, male: 51%, Fig. 2a). These data imply that, contrary to common belief, people are in fact not very accurate at reading a partner’s romantic intentions. Moreover, Fig. 2b shows that participants’ impression of being liked was associated with participants’ attraction to their partner (*F*(1, 402) = 64.55, *p* < 0.0001, Table S1). Yet, in reality, there was no association between how much participants were attracted to their partner and how much the other was attracted to them (*F*(1, 402) = 0.135, *p* = 0.71, Fig. 2c). These results align with previous literature showing that humans are poor perceivers of a partner’s intended flirtation and that they often project their own intentions onto their partner (32).

**Figure 2.**
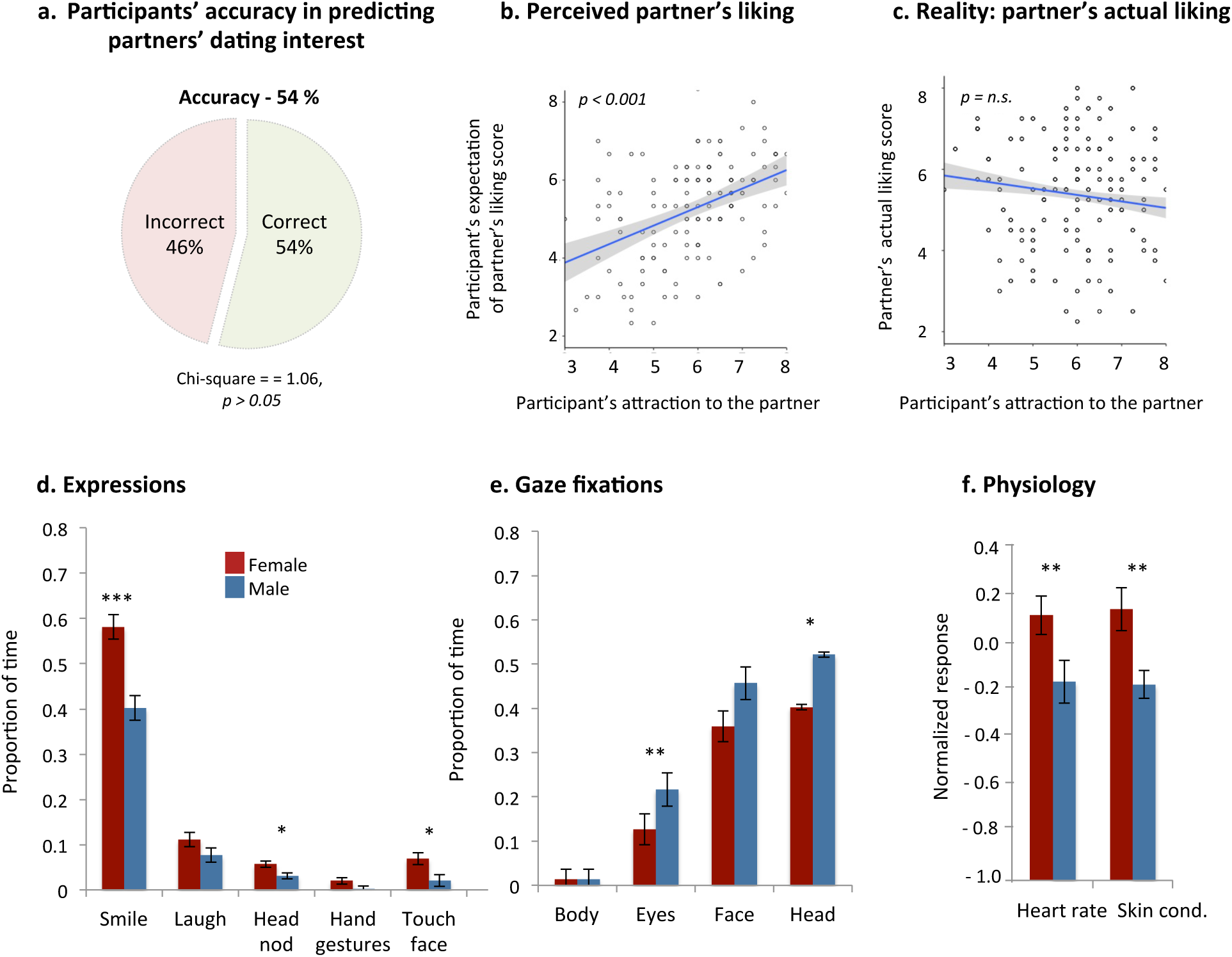
Behavioral and physiological results. **(a)** Pie chart shows the percentage of participants’ accuracy in predicting whether their partner wants to date them or not (binary decision, *N* = 138 couples): 74 people were accurate (54.4%), 62 people were inaccurate (45.6%); four people chose not to report. **(b)** Scatter plots show that the more males and females were attracted to their partner, the more likely they were to think that their partner was more attracted to them. **(c)** In reality, there was no association between how much participants were attracted to their partner and how much the other was attracted to them. Shaded areas represent 95% confidence intervals. Bar graphs represent gender differences in the proportion of time males and females displayed specific **(d)** expressions, **(e)** gazed at specific areas of interest and **(f)** average heart rate (HR) and skin conductance responses (SCR) across the three interaction types; physiological responses were normalized by baseline correction and z-transformation. Significance was defined using FDR 0.05. All \*\**p* < 0.01, \*\*\**p* < 0.001, *N =* 54 couples, error bars: ± SE.

### Individuals’ expressions

Having established that people are not good at predicting their partner’s romantic intentions, we set out to investigate whether participants’ facial expressions, eye fixations and physiological responses could be more reliable predictors of interpersonal attraction than participants’ judgment. We used two different approaches to answer this question. The first analysis was based on previous literature which suggests that behaviors such as smiling, direct eye contact and elevated arousal are signals that communicate romantic attraction, and therefore should be more evident when an individual is romantically or sexually motivated (33, 34). To find relevant patterns signaling attraction, first we tested whether males and females behave differently during the date (Table S2). The results obtained from a Multivariate generalized linear mixed model (F (11, 98) = 4.06, *p <* 0.0001; Pillai’s Trace = 0.34, Partial Eta = 0.34) indicated that females were significantly more expressive than males: females smiled, nodded and touched their face more frequently than males did (all *p*s < 0.01, Figure 2d). Males, on the other hand, stared at their female partner more; they fixated at females’ head and eyes significantly longer than females looked at them (all *p*s < 0.01, Fig. 2e), while females tended to look around and fixate longer at the background than males did (*p* = 0.025; Fig. S1). Additionally, females’ heart rate (F (1, 108) = 5.39, *p* = 0.002) and skin conductance responses (F (1, 108) = 9.68, *p* < 0.0001) were higher than males’ (Fig. 2f) and females also reported feeling more “aroused” and less self-confident than men (all *p*s < 0.01; Table S3-4). Together these data suggest that during a date, males’ and females’ behavior and physiology differs. Following this observation, we tested whether participants behaved in a specific way when they were attracted to their partners.

### Is there a specific behavioral or physiological pattern that predicts attraction?

As the results showed that males and females differ in frequency of expressions, gaze fixation duration and average physiological response, we split the data across genders and used machine-learning techniques to predict attraction. The major advantage of machine learning is that these methods can handle many predictors at the same time and, by using 10-fold cross-validation to validate our model, we can achieve an unbiased estimate of predictive performance of our models (35). In these models, we used participants’ expressions (i.e., frequency of smiling, laughing, hand gestures, head nods), eye fixations (e.g. duration of eye contact, face contact) and physiological responses (skin conductance, heart rate) as well as two-way interactions between all these features to predict males’ and females’ self-reported attraction scores during verbal and nonverbal interactions. Due to the high amount of predictors, we utilized a cross-validated Lasso model (36) to penalize non-predicted features (see methods). Intriguingly, the results showed that none of our observed *R*^*2*^s were significantly better than the permuted null distributions: men verbal interactions (*R*^*2*^ = −0.36, *p* = 0.96), men non-verbal interactions (*R*^*2*^ = −0.13, *p* = 0.48), women verbal interactions (*R*^*2*^ = −1.51, *p* = 0.92), women non-verbal interactions (*R*^*2*^ = −0.36, *p* = 0.96). This implies that attraction is not predictable by the participant’s autonomic nervous system activity, nor predicted by a specific frequency of expression or eye fixation duration. In other words, males and females who reported to be more attracted to their partner did not differ from those who were less attracted to their partner in their expressions, gaze fixations and physiological patterns. The same results were obtained from control GLM analyses, which also accounted for the multilevel and the longitudinal nature of our data (Table S5). In consequence, similarly to our participants (Fig. 2a), we were not able to accurately predict attraction level. In conclusion, the current data violates the first hypothesis suggesting that there are behavioral and physiological pattern that directly reflect feelings of attraction. To test the second hypothesis, we zoomed out from the individual and look at the couple as a whole.

### Couple’s expressions

In the second approach, we measured how synchrony between couples’ expressions, gaze and physiology affects participants’ attraction. To look for evidence of synchrony, we first used series of Spearman’s rank – order correlations and adjusted significance according to FDR Benjamini-Hochberg’s *p*-value (37). The results confirmed that, within couples, there were significant associations between the frequency of partners’ smiles, laughs, head nods, hand gestures and face touching (all ρ > 0.28, *p* < 0.001). Furthermore, participants also reciprocated each other’s gaze fixations as demonstrated by significant correlations between the duration that individuals looked eye-to-eye and head-to-head (all ρ > 0.22, *p* < 0.001). Finally, within the couples, we observed correlations in partners’ (baseline-corrected) average heart rate (ρ = 0.32, *p* < 0.001) and skin conductance levels (ρ = 0.36, *p* < 0.001). As a control analysis, we paired each female with a random male who did not belong to the real couple. In contrast to real couples, in randomly coupled participants, we did not find significant correlations in any of these measures (Fig. 2). An additional control analysis confirmed that the correlations between the shuffled dyads were significantly lower than the correlations in real dyads (all Fisher’s z > 0.2, *p* < 0.05, Table S6). Together these results demonstrate that participants synchronize with their partner on multiple levels of expressions, including gazing and facial expressions and physiology (Fig. 3a).

**Figure 3.**
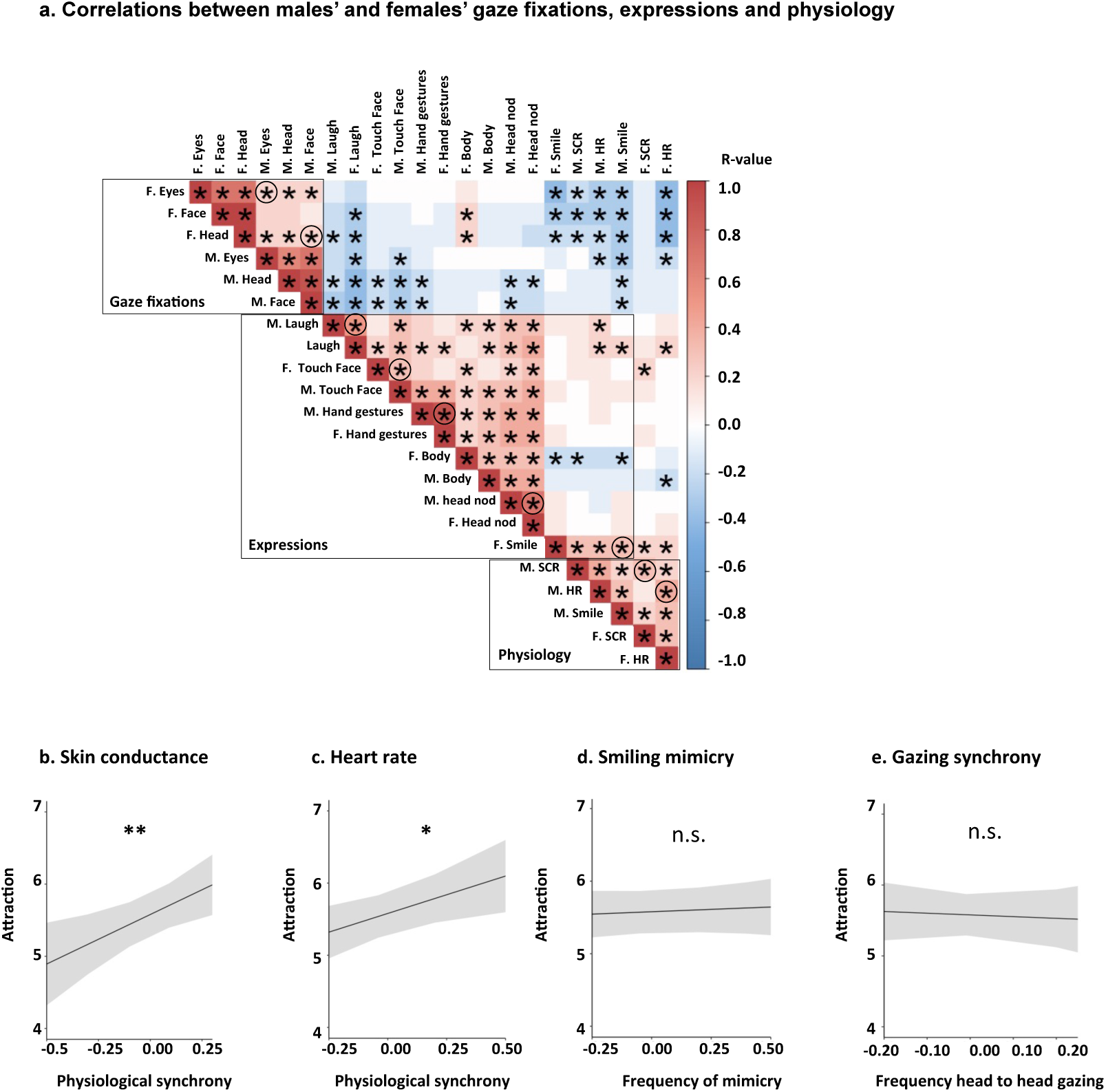
Synchrony and mimicry results. **(a)** Figure 3a displays correlation table summarizing associations between partners’ expressions, eye fixation and physiology for three interaction types (based on Spearman’s rank – order correlations, *N* = 162). F = females, M = males. The columns of the correlation matrix are placed according to the hierarchical clustering with similar values near each other. The black boxes framed around naturally occurring clusters demonstrate that synchrony occurred on all three levels of expressions; the circles represent significant mimicry between males’ and females’ eye-to-eye gaze, and head-to-head gaze. The circles in the physiology cluster highlight the significant relationship between males’ and females’ mean heart rate and skin conductance responses. The expressions’ circles signify mimicry between males’ and females’ smiling, hand gestures, head nods and laughter. The significance is adjusted according to FDR Benjamini-Hochberg’s *p*-value (37): \**p* < 0.05. **(b)** Attraction increase based on the synchrony of skin conductance levels [β = 3.044, SE = 0.95, CI (1.16, 4.93), *p* = 0.002] and **(c)** Attraction increase following heart rate synchrony during verbal interaction [*β* = 2.51, SE = 1.01, CI (0.52, 4.48), *p* = 0.013]. * *p* < 0.05, ** *p* < 0.01**. (d - e)** The frequency of smile mimicry and gaze synchrony did not significantly affect attraction. The shaded areas represent 95% confidence intervals. All predictors were centered. HR = heart rate, SCR = skin conductance response.

### Does synchrony promote attraction?

In the next analysis, we used Generalized linear mixed models to investigate how different types of interpersonal synchronies impact on participant’s attraction. In the initial model we included all 11 synchrony predictors including different types of expression mimicry (smiling, laughing, head shaking, hand gestures, face touching), gaze fixations synchrony (looking at partners’ head, eyes, face, body) and physiological synchrony (skin conductance, heart rate synchrony). The full model further included factors of gender, the type of interaction (verbal, nonverbal), the order of interaction (verbal/nonverbal first) and two-way interactions between all predictors as additional fixed effects. The final model was selected with a backward stepwise selection of fixed effects (Table S7).

Intriguingly, the final model showed that mimicry of visible signals that can be controlled, such as facial expressions or gaze, did not predict attraction (all *p*s > 0.05). On the other hand, attraction was predicted by synchrony in heart rate and skin conductance between partners, which is unconscious and difficult to regulate. Figure 3b-c shows that the more couples’ skin conductance levels synchronized (F (9, 314) = 8.87, *p* = 0.002) and the more heart rate responses synchronized during the verbal interaction (heart rate * interaction type: F (9, 314) = 6.21, *p* = 0.013), the more attracted couples became to each other over the course of the date. Crucially, we did not find this association in visible synchrony such as expression mimicry or alignment in gaze fixations (Figure 3d-e), which suggest that physiological synchrony explains more variance in attraction than the mimicry of explicitly visible expressions.

To test it the possibility that partners’ expressions promote attraction, regardless of synchrony. We conducted a follow-up control analysis (see Methods) in which instead of looking on participants’ attraction score we tested whether specific behavioral or physiological pattern enacted by one participant promotes attraction in the partner participant. The results of the control analyses revealed that people who were rated as attractive had no distinct expression pattern then those participants that were rates as less attractive (all *p*s > 0.05, full models summarized in Table S8). This finding supports the hypothesis that synchrony shapes social perception beyond individuals’ expressions (physiological or nonverbal). To show an example of what physiological synchrony looks like, we included a video of one couple (see Video 1). We selected this video because these two people first met without exchanging any words and, during this non-verbal interaction, their mean attraction score increased.

**Video 1. An example of physiological synchrony.** The video shows a nonverbal interaction where participants were instructed not to talk. At 00:04:00, the female will smile and the male partner reciprocates with a smile back. During this moment, we observe an increase in female’s and males’ skin conductance and heart rate (top two rows). Again, at 00:18:24, the female laughs; in response the male smiles and we again observe synchrony in heart rate and skin conductance. Thus, the purpose of the video is to explain how synchrony can occur. Importantly, not all smiles and laughs were paired with physiological synchrony, but in the case of this couple, they did. Further examination of these empirical visualizations suggested that physiological synchrony is more closely linked to “genuine” emotional exchange such as contagious smiles or uncontrolled laughter, as opposed to overt expressions used during polite communication (grins or nods).

## Discussion

In this study, we applied state of the art technology to define the processes that underlie human attraction in a real-life setting. First of all, we observed that people are not accurate at predicting their partner’s romantic intentions, as at the end of the date subjects’ predictions were at chance level. We further observed that participants who perceived their partner as highly attractive predicted that their partner liked them more than participants who rated their partner as less attractive. These results contradict the popular notion that people excel in their ‘mindreading’ capacities (38). Instead, our data suggest that during dating interactions people tend to mix their own feelings with partner’s feelings (33, 39).

Thanks to the combination of multiple measures we were able to acquire a new point of view, providing a more holistic understanding of nonverbal signals that drive social interactions. Specifically, in terms of nonverbal visible signals, we observed that females were more expressive than males during the date, which corresponds to previous research (40). Males, on the other hand, held their eye gaze more steadily focused on their partner than females. In addition, females were more physiologically and psychologically aroused compare to males. These are all novel findings, which we recommend replicating, as they provide fresh insights into the human sexual behavior. Furthermore, in line with prior literature (16, 20, 21, 28, 41–43), we found evidence for mimicry on all three levels of expression (expressions, eye gaze and physiology). Having established that people display variety of expressions, we set out to investigate whether participants’ facial expressions, eye fixations, and physiological responses could be more reliable predictors of interpersonal attraction. We used two different analytical approaches to tackle this question. The first approach was based on the hypothesis that individuals’ emotional expressions direct expression of an internal state, implying that they are inherently honest. Contrary to this prediction, results revealed that participants’ attraction could not be predicted by any of the visible signals (eye gaze, expression) nor subjects’ physiological responses (baseline-corrected HR, SCR). This result implicates that participants (males or females) who reported to be very attracted to their partner did not show any significant difference in their behavior or physiology than to those who were less attracted to their partner.

Instead, in support of the second hypothesis, we found that attraction was predicted by physiological synchrony between partners. Specifically, the level of skin conductance synchrony positively predicted attraction during both verbal and nonverbal interaction, and the level of heart rate synchrony predicted attraction during verbal interaction. Interestingly, attraction was not predicted by mimicry of visible expressions such as smiling, laughing, eye gazing (e.g., eye-to-eye contact), which implies that physiological synchrony captures more information than visible signals can explain. A plausible explanation of this finding is that visible social signals such as smile or eye-gaze can be easily controlled (12, 44). Since heart rate and skin conductance are autonomic signals that are difficult to regulate, physiological synchrony potentially signifies more ‘genuine emotional exchange’. In line with this interpretation, our data demonstrate that couples were often smiling and mimicking each other on a superficial level, yet these types of visible signals did not always align with people’s physiology. However, when participants’ physiological signals aligned during these interactions, attraction increased (Fig. 3). Even though we did not find strong evidence for the involvement of visible cues in attraction, this does not mean that overt expressions are not involved at all. In fact, video 1 depicts an example of a situation where physiological synchrony is triggered by nonverbal signals that couples dynamically exchange. Still, these visible signals and even their mimicry did not predict participants’ attraction ratings to the same extend as physiological synchrony did (Fig. 3). This is important as it suggest that physiological synchrony explains more information about interpersonal attraction than visible signals can convey. Although, the exact mechanisms through which autonomic synchrony affect social perception is not fully understood (20, 28). We propose that visible signals bring interacting partners’ autonomic activity into mutual alignment, creating a joint state that, which in turn encourages feeling of connection and mutual attraction.

The current findings are particularly relevant from the perspective of previous literature which suggest that behaviors such as smiling, direct eye contact and elevated arousal are signals that communicate romantic attraction (33, 34). In contrast, our result complements a recent meta-analytic review (12), which reported only a week relationship between self-reported attraction and affiliative behavior such as eye contact, smiling and mimicry. Most likely this is because during initial interactions people use emotional expressions to moderate social interaction, not necessarily to reveal their underlying subjective state. In the future, the physiology of synchrony might be studied in other social domains where nonverbal communication is the key. For instance, since autonomic cues are not likely to be influenced by learning, social norms, or conscious control (compared with facial expressions and other overt affective signals) the current set-up might be used to detect the onset of social deficits (e.g., bonding deficits or social anxiety). Furthermore, given that the current study is correlational, we recommend that further research tries to manipulate physiological synchrony to determine the causal link between synchrony and relationship formation.

In sum, thanks to the unique combination of measures (videos, eye-tracking and physiological measures), we were able to visualize the contagious spread of emotional information that stimulates attraction during real-life interactions. While in the field of social neuroscience, researchers have been mainly focusing on controllable expressions such as facial expression, body postures, and eye gaze (23, 38, 45); we show that visible signals or even their mimicry did not accurately predict the feeling of attraction. Instead, attraction was promoted by unconscious physiological synchrony between people. In conclusion, the current experiment goes beyond previous studies by linking the engagement of nonverbal mechanisms to physiological synchrony and attraction in real-life setting.

## Materials and Methods

### Participants

Our sample size was motivated by those used in previous studies (19, 46, 47). In total, 140 participants were recruited (70 opposite-sex dyads). Participants’ age ranged from 18 to 37 years old (Male: M = 25.71, SD = 4.639; Female: M = 23.45, SD = 4.265). Participants were recruited at three different yearly events in the Netherlands: during Lowlands (a music festival that takes place in the city of Biddinghuizen), The Night of Arts and Science (a festival that brings art and science together in Leiden) and during InScience (a science film festival in Nijmegen). To participate in the experiment, participants had to be single, between 18 and 37 years old, have normal vision or vision corrected by contact lenses (normal glasses could not be worn underneath the eye tracking glasses). Furthermore, participants could not have or have had any psychological illness, use medication or be undergoing psychological treatment. Using a digital 1PC alcohol tester we made sure to only include participants who did not exceed a blood alcohol content of 220 micrograms of alcohol per liter of exhaled breath (Dutch driving limit). For the behavioral analysis, one dyad was excluded because male left experiment prematurely; meaning 69 dyads were included in the behavioral analysis. For the physiological analysis an additional 15 dyads were excluded due to artifacts or missing physiological data, meaning 54 dyads were included in the physiological analysis. Participants were mostly Dutch (92%), and highly educated. Seventy-three percent of the subjects used dating applications (e.g., Tinder, Bumble, Happen) both males and females were looking for a committed relationship (see Table S9). At the end of the study, out of 138 people, in total 58 people (44%) wanted to date their partner at the end of the date (34% females, 53% males) from which eleven couples matched (17%), five people did not report. Furthermore, twenty couples (31%) mutually agreed on not being a good match for each other and in half of the couples (52%) one partner wanted to date their partner but the other did not reciprocate. There were no significant differences between males and females in their level of social anxiety, positive/negative affect or score on the social desire scale (Table S10). The experimental procedures were in accordance with the Declaration of Helsinki and approved by the Ethical Committee of the Faculty of Behavioral and Social Sciences of the University of Amsterdam. All participants provided informed consent.

### Procedure

#### Baseline measures

Participants were screened for exclusion criteria, received information about the study and gave informed written consent. Subjects were then asked to fill out some control questionnaires to control for psychological factors that could influence a person’s ratings of their partner or the general behavior during social interactions (see Materials). In addition, participants filled out baseline ratings reporting on their expectations and standards (e.g. how attractive, intelligent, trustworthy and funny their potential romantic partner should be). Subjects also rated themselves on the same items on the 10-point scales.

Two researchers (one for male, one for female) attached electrodes measuring heart rate (HR) and skin conductance (SC) to participants’ skin. They also helped participants to put on the eye-tracking glasses, which were calibrated afterward. Without seeing their partner, participants were led to the dating cabin, females first and after calibration of her equipment, the male partner followed. Upon eye-tracking and skin conductance calibration, participants were instructed to look at the fixation cross (at the closed barrier), while their baseline (30 seconds) physiological measures were collected. Cameras in the glasses recorded video and sound over the whole period of the dating experiment. Participants were instructed to remain silent until they heard instructions via a speaker.

#### First impression

The screen then opened shortly (3 seconds), giving participants a first impression of their partner. After the first impression, participants looked at the fixation cross for 30 seconds to collect post-first impression physiological measures after which they rated their partner on the same (0 – 9) scales as they rated their imaginary or potential romantic partner during baseline. In addition, participants were asked to rate how much they liked their partner and how much they thought their partner liked them. Other questions included how similar they thought the partner was in terms of personality and how much connection, ‘click’, and sexual attraction they felt between them. After the first impression, two additional interactions would take place (the order of which was counterbalanced).

#### Verbal interaction

The visual barrier opened and participants were instructed to talk freely with their partner for 2 minutes. After this interaction, the participant was asked to fill in the same scales as during the first impression, plus rate their impression of the verbal interaction.

#### Nonverbal Interaction

The visual barrier opened and participants were instructed to look at their partner and not speak for 2 minutes. Afterward, the barrier closed and subjects rated their partner on the same 0 – 9 point scales. Whether participants began with verbal or nonverbal interactions was counterbalanced (Fig. 1b). During the final ratings, participants indicated how much they thought the other person liked them and whether they wanted the experimenters to exchange their email addresses. The pairs were also asked to predict whether they thought their partner wanted to exchange email and go for another date. Finally, subjects were asked to indicate whether their video recordings could be used for follow-up experiments.

#### Follow-up

For ethical reasons, participants’ decisions to date their partner again or not were not revealed until the festival was over. Only if both of them agreed to exchange contact information, one week after the study they have received an email with their partner’s email address. They were asked if we could contact them again later to ask if they were still in contact with their partner.

### Measures

#### Ratings

Participants filled in ratings before the experiment, after the first impression and after both the verbal and nonverbal interactions. All questionnaires included the same questions about the partner (or during baseline about a potential partner) in which the participant rated: attraction, funniness, intelligence, trustworthiness, the similarity in personality, connection, sexual attraction and click, on scales ranging from 1 (not at all) to 9 (very). Additionally, during baseline, participants had to indicate how attractive, funny, intelligent and trustworthy they thought they themselves were (0 – 9 scales). Every questionnaire also contained a mood grid, in which participants had to indicate their level of arousal and valence of their affect. Subjects also rated how shy, awkward and self-confident they were feeling. Furthermore, every questionnaire (except during baseline), included a question asking how much they liked the partner, and how much they thought their partner liked them. Finally, during the first impression and during their last interaction, participants indicated whether they wanted to see their partner again and whether they thought their partner wanted to see them again. As additional control measures, we included the Liebowitz Social Anxiety Scale (48), Positive and Negative Affect Schedule (49) and Sexual Desire Inventory (50).

### Pre-processing

#### Behavioral expressions coding

The eye-tracking glasses automatically detected eye-fixations and videotaped participants’ behavior. Four independent raters (two raters for males and two for females) rated participants’ expressions (smiling, laughing, head nod, hand gestures, face touching) using the Tobii Pro Lab (Version 1.5, 5884). The tapes were coded without sound and coders were blind to participants’ ratings. The facial expressions were coded per tenths of seconds and the frequency of each expression was then averaged per interaction (lasting between 3 seconds – 120 seconds). The reliability then was calculated as percentage of agreement between recoded observations. All coders had successfully completed training and reached an agreement ratio of at least .70 for all behaviors, except for the open versus closed body position (agreement was less than 70%); thus this particular behavior was dropped from all analyses.

#### Eye gaze fixations classification

Eye fixations were recorded using Tobii Pro Glasses 2. We defined areas of interest (AOI) including the head, face, eyes, nose, mouth, body, right arm, left arm and background. AOIs were drawn on snapshot images of participants taken at the start of each interaction (size in pixels: 1079 x 605). Eye gaze fixations were then automatically mapped onto the areas of interest (partner’s face and body) using the Fixation Classification Method implemented in Tobii Pro Lab (Version 1.5, 5884). The I-VT (Attention) filter (Velocity-Threshold Identification Gaze Filter) was selected to handle eye-tracking data from glasses recordings conducted under dynamic situations. Like with expressions, the fixations were collected per tenths of seconds for each AOI. This resulted in AOI visit duration (0 excluded). Prior to each interaction, we checked whether the eye-tracker needed recalibration or not. To do so, we asked participants to focus on the fixation point at the barrier. In case the eye fixation did not overlay the fixation cross, we re-calibrated. In the post-experiment pre-processing stage, we calculated the remaining small differences in the x and y coordinates between the glasses fixation and the fixation cross. The AOI masks were moved with the small differences on the respective x and y coordinates.

#### Physiological measures

For each participant, ECG and EDA data were collected using BIOPAC’s ECG2-R and PPGED-R modules, respectively, and an MP-150 system operated using AcqKnowledge software version 3.2 (BIOPAC, Goleta, CA). All raw signals were recorded at 1000 Hz.

#### Skin conductance pre-processing

Using the PhysioData Toolbox, the raw skin conductance signal was visually inspected and short-duration artifacts were removed and replaced using linear interpolation. Longer invalid sections of data were excluded. The skin conductance signal (SC) was then low-pass filtered at 2 Hz to remove high-frequency noise, and for each section of interest, down-sampled to 10 Hz for further analysis.

#### Heart rate pre-processing

Similarly, the PhysioData Toolbox was used to extract 10 Hz continuous instantaneous heartrate (IHR) signals from the raw ECG signal. This involved bandpass-filtering the raw signal at 1 to 50 Hz, performing peak detection to find the R-peaks, and calculating the interbeat intervals (IBIs). Both the R-peaks and resulting IBIs were visually reviewed, and erroneously derived instances of any of the two were removed. The IHR signal, in BPM, was then generated from the remaining IBIs using piece-wise cubic interpolation. Sections missing more 50% percent of the IBIs were excluded.

### Analysis

#### Analysis 1

At the final ratings, we asked participants whether they thought their partner wanted to date them or not (yes =1/no=2). We subtracted these answers from partners’ actual response (partner really wants to date: yes =1/no=2). This resulted in either accurate (0), false negative (1) or false positive (−1) answers. We then binarized the accuracy variable (correct/incorrect (pooling 1 and −1)) to test for significance above chance level with Chi-square test (alpha = 0.05).

#### Analysis 2

Apart from categorical answers (date or not), we asked participants to rate how attractive their partner was and how much they thought partner liked them. We asked these questions to investigate whether participants’ impression of being liked related to their attraction towards their partner. We used a Multilevel linear mixed model with 3 level structure: dyad (Level 3), participant (Level 2) and time (Level 1). In this model, participants’ impression of being liked was used as the target variable predicted by participant’s attraction towards their partner. We further included gender and the interaction between gender and attraction to control for gender differences (see Table S1).

#### Analysis 3

We tested whether females and males differ in frequency of their naturally occurring expressions, eye fixations, and physiological responses. To do so, for each interaction type (first impression, verbal interaction and nonverbal interaction) and each participant we calculated the proportion of time (min = 0, max = 1) that participants were (i) smiling, (ii) laughing, (iii) head shaking, (iv) making hand gestures, (v) touching their face or fixating on partners’ (vi) body, (vii) eyes, (viii) face, (ix) head. Physiological levels: (x) skin conductance and (xi) heart rate responses were baseline corrected (30 seconds prior to every interaction) and then z-scored. This resulted in eleven averaged values for each subject and interaction. We used a 3 x 2 Multivariate Generalized Linear Mixed model to test for gender differences using the within subject factor interaction type (first impression, verbal and nonverbal interaction), gender (male, female) as between subject factor. To control for multiple comparisons we employed a false discovery rate (FDR) in all following models (37). To check whether females look longer at the background than males do, we conducted a Generalized Linear Mixed Model. In this model, the data were nested in each subject and the individual intercept was random. The average time (in seconds) looking at the background was used as a dependent variable and gender, interaction type (first impression, verbal, nonverbal), gender * interaction type were used as fixed effects (see Fig.1).

#### Analysis 4

Apart from physiological arousal, we investigated whether males and females differ in their cognitive arousal by conducting Multivariate Generalized Linear Mixed model testing for gender differences on mood grids: (i) arousal (ii) valence, self-ratings reporting the level of (ii) shyness and (v) self-confidence.

#### Analysis 5

Furthermore, we tested whether males and females behaved in a specific way when they felt attracted to their partner. We used machine-learning techniques to detect a specific behavioral and physiological pattern that would predict participant’s attraction level. Using a 10-fold cross-validated lasso regression model (alpha = 0.5) implemented in Sci-kit learn in python 3.6 (51), we aimed to predict attraction directly following either the verbal or non-verbal interaction, separately for men and women. In our model, we included all the predictors including participants’ frequency of smiling, laughing, hand gestures, fixations duration on partner’s eyes, face, head, body and physiological responses (skin conductance, heart rate) as well as 2-way interactions between all those features (91 predictors in total) to predict males’ and females’ self-reported attraction scores. All predictors were z-scored. To evaluate the performance of our models we performed a permutation test with 3000 permutations, shuffling the attraction levels across participants and testing that our observed non-shuffled R^2^ was larger than 95% of the randomly shuffled R^2^.

#### Control Analyses 5

To account for within subject and dyad dependencies, we conducted a series of Multilevel mixed effects models with following structure: three time points (Level 1) nested in participants (Level 2), nested in dyads (Level 3). In each model, expression, gaze fixations frequencies and baseline corrected physiological responses were used as predictors of participant’s attraction scores (scale 0 – 9). Gender and gender * expression/fixation interaction were used as additional predictors of attraction. Due to the multicollinear nature of the data, we carried out a model for each expression, gaze fixations frequencies and physiological response independently (11 mixed effects models). The Multilevel mixed effects models were conducted such that the intercept terms were allowed to vary across dyads and participants, we further used an AR1 covariance matrix to account for time dependencies. We defined significance using an FDR < 0.05. The results of control analyses complimented the results of Analysis 5 (see Tables 5-7).

#### Analysis 6

We ran a correlation between all measures. This resulted in a large correlation table showing associations between male’s and female’s expressions eye fixations and physiological measures as well as associations between female’s-female’s, male’s-male’s showing how nonverbal behaviors and physiological responses relate to each other within participants. Then in control analysis, each female was paired with a random male. To test for significance, we directly contrasted the (FDR corrected) correlations coefficients between true couples and randomly matched couples with cocor package in R studio (52) using gender an independent group, two sided test with alpha set to 0.05.

#### Quantifying expressive mimicry and eye fixation synchrony

We quantified mimicry for each dyad and interaction by calculating the proportion of time both participants’ directly reciprocated expressions (smiling, laughing, head nod, hand gestures, face touching) and gaze fixations (looking at partners’ head, eyes, face, body).

#### Quantifying synchrony

We quantified synchrony with windowed cross-lagged correlational analyses (31). This method has the advantage that it takes into account the non-stationarity of the time series and the dynamical nature of the interaction. This is important as the level of synchrony may fluctuate during the experiment. The first step in the analysis was to determine the parameters (window size, window increment, maximum lag, lag increment). We did that following an extensive process by comparing previous studies using similar statistical methods, looking at what is physiologically plausible given the time course of the physiological signals and by employing a data-driven bottom-up approach where we investigated how changing the parameters affected the outcomes using a different dataset. As expected, the absolute values of the synchrony measures varied depending on the parameters (e.g., the window size, the lag size), but as supported by McAssey, Helm, Hsieh, Sbarra, and Ferrer (53), the relative results were not affected (e.g. a dyadic manifesting relatively high synchrony showed such tendency for the different parameters). Based on these three factors, we set the parameters as follows: the window size was 8 seconds, the window increment was 2 seconds, the maximum lag was 4 seconds and the lag increment was 100ms. Then a peak picking algorithm was applied (31). This algorithm allows for detecting the maximum cross-correlation across the lags for each time segment. Both the windowed cross-correlations and the peak picking algorithm were conducted 6 times per dyad, once for the heart rate responses and once for the skin conductance responses for each condition (the first impression, verbal and nonverbal interaction) resulting in N dyads * 6 result and peak picking matrices. Finally, the mean and standard deviation of the peak cross-correlations of all window segments and the mean of the absolute values of corresponding time lags were calculated for both physiological measures for each condition per dyad.

#### Analysis 7

We test whether attraction can be predicted by synchrony. In this model, we used synchrony in expressions (smiling, laughing, head nod, hand gestures, face touching) and gaze fixations (looking at partners’ body, head, eyes, face) and physiology (skin conductance, heart rate) as predictors of participant’s attraction. In addition, gender, interaction type (verbal, nonverbal), the order of interaction (verbal/nonverbal first) were used as additional predictors in the model. To allow for differences between dyads, the intercept terms were allowed to vary across dyads and we included a first-order autoregressive AR(1) residuals structure to account for time dependencies. The final model was selected with a backward stepwise selection of fixed effects in a generalized linear mixed-effects model. This method first tests interaction terms, and then drops interactions one by one to test for main effects. Main effects that are part of interaction terms were retained, regardless of their significance as main effects (full model summarized in Table 9).

#### Control analyses 7

We conducted a follow-up control analyses, in which similarly to analyses 5 and 7, we used cross-validated Lasso model and generalized linear mixed-effects model (same structure as in analysis 7) to predict whether participants’ expressions (i.e., frequency of smiling, laughing, hand gestures, head nods), eye fixations (e.g. duration of eye contact, face contact) and physiological responses (skin conductance, heart rate) as well as two-way interactions between features predict self-reported attraction scores. This time, instead of looking on participants’ attraction score we tested whether specific behavioral or physiological pattern enacted by one individual promotes attraction in the other individual. The results of control analysis showed that people who are rated as attractive have no distinct expression pattern then those participants that were rates as less attractive.

## Data and code availability

All data, code, and materials that are associated with this paper and used to conduct the analyses are accessible at: https://github.com/Eprochazkova/Physio_Synchrony

## Acknowledgments

We thank Michael Rojeck Giffin for helpful feedback, Elio Sjak-Shie for technical assistance, and Wouter Boekel for proof-reading the scripts. Research was supported by the Netherlands Science Foundation (VENI # 016-155-082) to Mariska E. Kret and (Talent Grant # 406-15-026) from NWO (Nederlandse Organisatie voor Wetenschappelijk Onderzoek) to Mariska E. Kret and Eliska Prochazkova.

## Author contributions

E.P. conceived the idea. E.P, M.E.K. and F.B. designed experiment and with contributions from D. L. conducted the experiment. E.P., E.S.S and D.L. performed the analyses and computational modeling with contributions from M.E.K wrote the paper with contributions from F.B. All authors discussed the results and implications and commented on the manuscript at all stages.

## Competing interests

The authors declare no competing financial interests.

## Materials & Correspondence

Correspondence and material requests should be addressed to E.P and M.E.K

