## Supplementary Materials for "The choreography of human attraction: physiological synchrony in a blind date setting"

**This PDF file includes:**

Figures S1 to S2

Tables S1 to S10

Legends for Movies S1

**Other supplementary materials for this manuscript include the following:**

Movies S1

Supplementary Figures

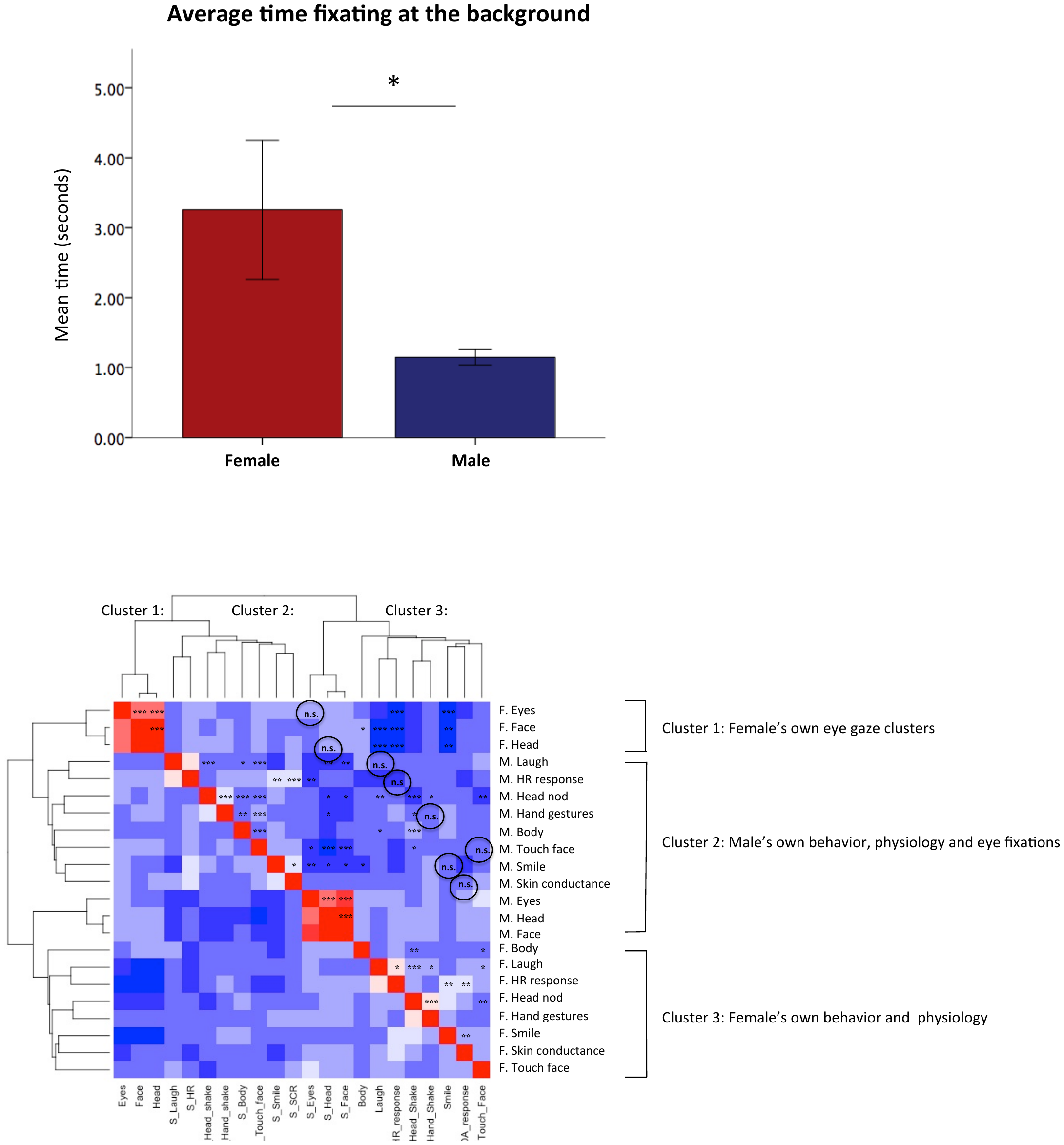

**Fig. S1.** **Control analysis:** Bar plot shows mean time males and females fixated at the background during interactions. Error bars: ± SE.

To check whether females look longer at the background than males do, we conducted a Generalized Linear Mixed Model. In this model, the data were nested in each subject, the average time looking at the background (in seconds) was used as the dependent variable and gender, interaction type (First impression, verbal, nonverbal), gender * interaction type were used as fixed effect. The intercepts were allowed to be random across participants. The results confirmed the main effect of gender (F(1, 347) = 2.637, *p* = 0.025), as females gazed at the background on average 2 seconds longer than males did (β = 2.104, SE = 0.93, CI (0.26, 3.94), *p* = 0.025). There was no significant difference between the interaction types (*p* < 0.05) and the interaction between gender*interaction type (*p* < 0.05).

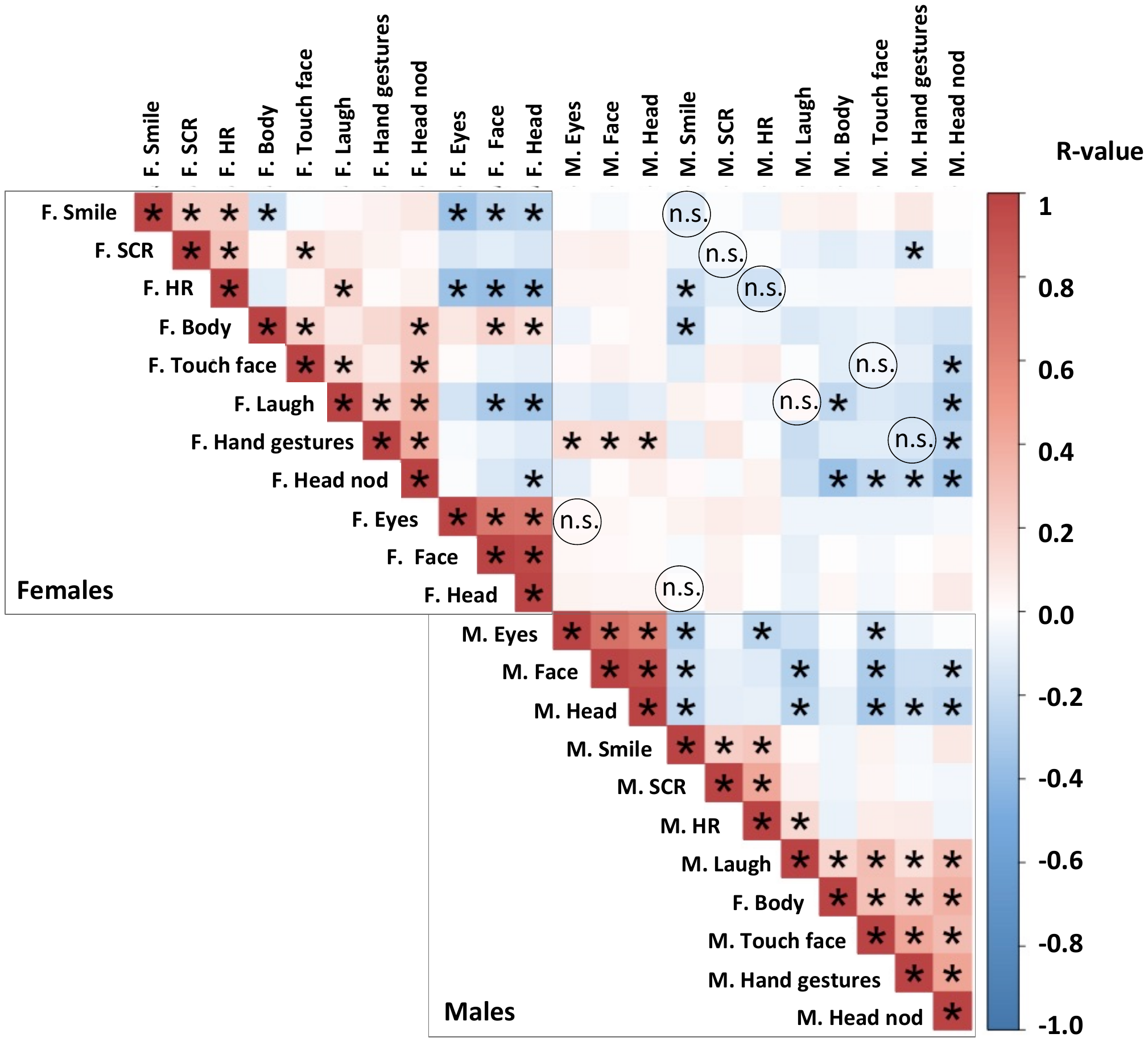

**Fig. S2. Correlation table summarizes** associations between randomly matched males and females and within-subject correlations in participants’ expressions, fixations and physiology. F. = real female, M. = randomly paired male. The heat map shows that in randomly paired couples the associated were formed only within subjects, while in real couples the behavior clustered also between male and female participants, we used the FDR < 0.05 to define significance, (N = 162).

Supplementary Tables

**Table S1a.** Attraction as a predictor of the participants’ impression of being liked

| (IV) | F | Df1, Df2 | *P*-value | Adjusted *P* value | FDR Significant |
| --- | --- | --- | --- | --- | --- |
| Attraction (intercept) | 22.63 | 3, 402 | 0.00 | 0.00 | Yes |
| Attraction | 64.54 | 1, 402 | 0.00 | 0.00 | Yes |
| Gender | 0.14 | 1, 402 | 0.71 | 0.71 | No |
| Gender *Attraction | 0.78 | 1, 402 | 0.60 | 0.80 | No |

Covariance Parameters

| Residual effects | Est. | Std. Er | Z | *P*-value | 95% CI: lower | upper |
| --- | --- | --- | --- | --- | --- | --- |
| Intercept (dyad) | 0.06 | 0.18 | 0.33 | 0.743 | 0.00 | 0.88 |
| Smile (AR1 Rho) | 1.014 | 0.23 | 4.32 | 0.00 | 0.64 | 1.59 |

**Table S1**: The results of multilevel linear mixed model with 3 level structure: dyad (Level 3), participant (Level 2) and time (Level 1). In this model, Participants’ impression of being liked was used as the target variable predicted by participant’s attraction towards their partner. We further included gender and the interaction between gender and attraction and FDR < 0.05 to define significance.

**Table S2.** Gender differences in expression frequency, gaze frequency and physiological responses

| Dependent Variable | Type III Sum of Squares | Mean Square | F | *P*-value | Partial Eta Squared | Noncent. Parameter | Obs. Power | Adjusted *P* value | FDR Significant |
| --- | --- | --- | --- | --- | --- | --- | --- | --- | --- |
| Fixations on Eyes | 0.61 | 0.61 | 7.86 | 0.01 | 0.04 | 12.79 | 0.95 | 0.02 | Yes |
| Fixations on Face | 0.70 | 0.70 | 3.51 | 0.06 | 0.02 | 7.01 | 0.75 | 0.09 | No |
| Fixations on Head | 1.06 | 1.06 | 5.61 | 0.02 | 0.04 | 11.28 | 0.92 | 0.04 | Yes |
| Fixations on Body | 0.00 | 0.00 | 0.01 | 0.92 | 0.00 | 0.02 | 0.05 | 0.92 | No |
| Smile | 2.43 | 2.43 | 17.14 | 0.00 | 0.06 | 19.29 | 0.99 | 0.00 | Yes |
| Laugh | 0.06 | 0.06 | 0.97 | 0.33 | 0.00 | 1.19 | 0.19 | 0.36 | No |
| Head nod | 0.05 | 0.05 | 6.41 | 0.01 | 0.02 | 5.40 | 0.64 | 0.04 | Yes |
| Hand gestures | 0.03 | 0.03 | 1.16 | 0.29 | 0.01 | 3.22 | 0.43 | 0.35 | No |
| Touching Face | 0.20 | 0.20 | 5.02 | 0.03 | 0.02 | 7.26 | 0.77 | 0.04 | Yes |
| Mean HR response | 7.74 | 7.74 | 5.39 | 0.02 | 0.02 | 5.52 | 0.65 | 0.04 | Yes |
| Mean SCR response | 10.53 | 10.53 | 9.68 | 0.00 | 0.03 | 9.90 | 0.88 | 0.01 | Yes |

**Table S2** shows 3 x 2 mixed effect MANOVAs tested for gender differences using within subject factor interaction type (first impression, verbal and nonverbal interaction) and gender (male, female) as between subject factor. Significance was defined using FDR 0.05 (Benjamini-Hochberg Adjusted *P*-value). SCR: Skin conductance, HR: Heart rate

| **Table S3.** Descriptive statistics behavioural ratings between males and females | | | | | |
| --- | --- | --- | --- | --- | --- |
| Dependent Variable | Gender | Mean | Std. Error | 95% Confidence Interval | |
|  |  |  |  | Lower | Upper |
| Valence | female | 5.80 | 0.12 | 5.56 | 6.04 |
|  | male | 6.18 | 0.13 | 5.93 | 6.43 |
| Arousal | female | 6.20 | 0.13 | 5.95 | 6.46 |
|  | male | 5.71 | 0.14 | 5.44 | 5.98 |
| Shy | female | 4.50 | 0.15 | 4.21 | 4.79 |
|  | male | 4.18 | 0.15 | 3.87 | 4.48 |
| Awkward | female | 5.19 | 0.16 | 4.87 | 5.50 |
|  | male | 4.29 | 0.17 | 3.96 | 4.62 |
| Self confidence | female | 5.28 | 0.11 | 5.06 | 5.49 |
|  | male | 5.98 | 0.11 | 5.76 | 6.20 |

| **Table S4.** MANOVAs compare behavioural ratings between males and females | | | | | | | | |  |
| --- | --- | --- | --- | --- | --- | --- | --- | --- | --- |
| Dependent Variable | Sum of Squares | Mean Square | F | *P*-value | Partial Eta Squared | Noncent. Parameter | Observed Powerf | Adjusted *P* value | FDR Significant |
| Valence | 12.84 | 12.84 | 4.77 | 0.03 | 0.01 | 4.77 | 0.59 | 0.04 | Yes |
| Arousal | 21.14 | 21.14 | 7.03 | 0.01 | 0.02 | 7.03 | 0.75 | 0.01 | Yes |
| Shy | 9.10 | 9.10 | 2.32 | 0.13 | 0.01 | 2.32 | 0.33 | 0.13 | No |
| Awkward | 69.46 | 69.46 | 15.08 | 0.00 | 0.04 | 15.08 | 0.97 | 0.00 | Yes |
| Self confidence | 43.27 | 43.27 | 20.70 | 0.00 | 0.06 | 20.70 | 1.00 | 0.00 | Yes |

We used the FDR < 0.05 to define significance.

**Table 3 – 4** shows the results obtained from Multivariate GLMs comparing males and females ratings of emotional arousal, valence, shyness, awkwardness and self-confidence showed a main effect of gender (F (5, 341) = 8.252, *p* < 0.0001; Pillai’s Trace = 0.108). We used the FDR < 0.05 to define significance, (N = 162).

**Table S5.** **Expansions, Eye Fixations and Physiological response** as a predictor of the participants’ romantic attraction scores

| Expressions (IV) | F | Df1, Df2 | *P*-value | FDR Significant |
| --- | --- | --- | --- | --- |
| Gender | 8.62 | 1, 320 | 0.00 | Yes |
| Smile | 0.33 | 1, 320 | 0.57 | No |
| Laugh | 0.15 | 1, 320 | 0.70 | No |
| Hand gestures | 0.00 | 1, 320 | 0.90 | No |
| Head shake | 1.99 | 1, 320 | 0.16 | No |
| Touch Face | 0.91 | 1, 320 | 0.34 | No |
| Body | 0.69 | 1, 320 | 0.41 | No |
| Eyes | 0.63 | 1, 320 | 0.42 | No |
| Face | 0.34 | 1, 320 | 0.56 | No |
| Head | 0.00 | 1, 320 | 0.95 | No |
| HR | 0.06 | 1, 276 | 0.80 | No |
| SCR | 0.37 | 1, 239 | 0.54 | No |
| Smile*Gender | 0.34 | 1, 320 | 0.57 | No |
| Laugh*Gender | 2.11 | 1, 320 | 0.15 | No |
| Hand gestures*Gender | 0.00 | 1, 320 | 1.00 | No |
| Head shake*Gender | 0.04 | 1, 320 | 0.83 | No |
| Touch Face*Gender | 1.69 | 3, 320 | 0.40 | No |
| Body*Gender | 0.49 | 1, 320 | 0.48 | No |
| Eyes*Gender | 1.06 | 1, 320 | 0.42 | No |
| Face*Gender | 1.78 | 1, 320 | 0.18 | No |
| Head*Gender | 2.26 | 1, 320 | 0.13 | No |
| HR*Gender | 0.46 | 1, 276 | 0.49 | No |
| SCR*Gender | 0.94 | 1, 239 | 0.33 | No |

**Tables S5** summarize series of Generalized linear mixed models with attraction scores as the target (scale 0 – 9; the intercept varied across dyads and participants). Due to the multicoliniar nature of the data, we carried out 11 mixed effects models for each participant’s expression independently. In each model, expressions (smiling, laughing, head shaking, hand gestures, face touching) and gaze fixations (looking at partners’ head, eyes, face, body) and physiological baseline corrected response (skin conductance, heart rate) were used as predictors. Gender and the interaction between gender * expression were used as additional predictors. We defined significance using an FDR < 0.05. The results of Multilevel mixed effects models revealed that none of the directly visible signals such as participants’ expressions and gaze fixations were significant predictors of male’s or female’s self-reported attraction scores (all *p*s > 0.05, full models summarized in Supplementary Table 5-6). Also, contrary to common assumptions, even when we looked deeper into people’s physiology, participants’ heart rate or skin conductance levels during the date did not predict how attracted participants were to their partner (all *p*s > 0.05, Supplementary Table 7). This implies that attraction is neither linearly related to the participant’s autonomic nervous system activity, nor predicted by a specific frequency of expression or eye fixation duration.

**Table S6.** Comparisons between true couple and randomly matched couples

|  | True Male | Random Male | Fisher's Z | Significance |
| --- | --- | --- | --- | --- |
| Female's Eyes | 0.69 | 0.34 | 4.4 | 0.00 |
| Female's Head | 0.22 | -0.19 | 0.28 | 0.78 |
| Female's Face | 0.14 | 0.13 | 0.09 | 0.93 |
| Female's Skin conductance | 0.32 | -0.07 | 10.69 | 0.00 |
| Female's Heart rate | 0.36 | -0.07 | 11.09 | 0.00 |
| Female's Smile | 0.31 | 0.02 | 2.68 | 0.01 |
| Female's Laugh | 0.51 | 0.28 | 7.58 | 0.00 |
| Female's Touch Face | 0.27 | 0.25 | 0.19 | 0.85 |
| Female's Head nod | 0.66 | -0.23 | 9.16 | 0.00 |
| Female's Hand gestures | 0.87 | 0.03 | 9.13 | 0.00 |

**Table S6** displays Correlations between real couples and randomly matched couples were tested with cocor package in R studio using gender an independent group, alpha set to 0.05 (two sided test). Note that there is not a significant difference in touching face and face/head fixations between true couples and random couples. We defined significance using an FDR < 0.05.

**Table S7a.** The final multilevel model: synchrony predicts attraction

| Expressions (IV) | F | Df1, Df2 | P-value | Adjusted P value | FDR Significant |
| --- | --- | --- | --- | --- | --- |
| Corrected model (intercept) | 4.03 | 9, 314 | 0.00 | 0.00 | Yes |
| Interaction type | 0.01 | 1, 314 | 0.91 | 0.91 | No |
| Gender | 10.5 | 1, 314 | 0.00 | 0.01 | Yes |
| Order interactions | 3.14 | 1, 314 | 0.08 | 0.16 | No |
| Skin conductance synchrony | 8.73 | 1, 314 | 0.00 | 0.01 | Yes |
| Heart rate synchrony | 0.25 | 1, 314 | 0.61 | 0.68 | No |
| Gender*Heart rate synchrony | 0.27 | 1, 314 | 0.61 | 0.76 | No |
| Gender*Skin conductance synchrony | 2.14 | 1, 314 | 0.14 | 0.21 | No |
| Interaction type*Heart rate synchrony | 6.21 | 1, 314 | 0.01 | 0.03 | Yes |
| Interaction type*Skin conductance synchrony | 2.92 | 1, 314 | 0.09 | 0.15 | No |

**Table S7b.** Covariance Parameters

| Residual effects | Est. | Std. Er | Z | P-value | 95% CI: lower | upper |
| --- | --- | --- | --- | --- | --- | --- |
| Smile (Diagonal) | 2.75 | 0.30 | 9.29 | 0.00 | 2.23 | 3.40 |
| Smile (AR1 Rho) | 0.73 | 0.04 | 21.14 | 0.00 | 0.66 | 0.79 |

**Table S7a-7b** shows the final model selected with a backward stepwise selection of fixed effects in a generalized linear mixed-effects model. In the initial model we included all 11 synchrony predictors (smiling, laughing, head nod, hand gestures, face touching, looking at partners’ body, head, eyes, face, skin conductance, heart rate), gender, interaction type (verbal, nonverbal), the order of interaction (verbal/nonverbal first) and two-way interactions between all predictors as additional fixed effects. To allow for differences between dyads, the intercept terms were allowed to vary across dyads and we included a first-order autoregressive AR(1) residuals structure to account for time dependencies. The final model drops interactions one by one to test for main effects. We defined significance using an FDR < 0.05.

**Table S8.** **Control analysis:** Expansions, Eye Fixations and Physiological response as a predictor of the partner’ romantic attraction scores

| Expressions (IV) | F | Df1, Df2 | *P*-value | FDR Significant |
| --- | --- | --- | --- | --- |
| Gender | 8.59 | 1, 320 | 0.04 | Yes |
| Smile | 1.61 | 1, 320 | 0.21 | No |
| Laugh | 0.00 | 1, 320 | 0.98 | No |
| Hand gestures | 0.52 | 1, 320 | 0.47 | No |
| Head shake | 0.04 | 1, 320 | 0.85 | No |
| Touch Face | 0.00 | 1, 320 | 0.95 | No |
| Body | 0.65 | 1, 320 | 0.42 | No |
| Eyes | 0.09 | 1, 320 | 0.77 | No |
| Face | 1.41 | 1, 320 | 0.56 | No |
| Head | 1.81 | 1, 320 | 0.18 | No |
| HR | 0.29 | 1, 276 | 0.59 | No |
| SCR | 0.00 | 1, 239 | 0.99 | No |
| Smile*Gender | 0.94 | 1, 320 | 0.57 | No |
| Laugh*Gender | 1.93 | 1, 320 | 0.17 | No |
| Hand gestures*Gender | 0.00 | 1, 320 | 1.00 | No |
| Head shake*Gender | 0.00 | 1, 320 | 0.98 | No |
| Touch Face*Gender | 0.72 | 3, 320 | 0.40 | No |
| Body*Gender | 0.90 | 1, 320 | 0.48 | No |
| Eyes*Gender | 0.20 | 1, 320 | 0.66 | No |
| Face*Gender | 0.71 | 1, 320 | 0.40 | No |
| Head*Gender | 1.53 | 1, 320 | 0.22 | No |
| HR*Gender | 0.14 | 1, 320 | 0.71 | No |
| SCR*Gender | 0.08 | 3, 320 | 0.78 | No |

**Table S9: Participants’ demographics**

|  | Female | | |  | Male | | |  |
| --- | --- | --- | --- | --- | --- | --- | --- | --- |
| Characteristics | N | M | SD |  | N | M | SD | *t* |
| Age | 69.00 | 23.45 | 4.18 |  | 69.00 | 25.71 | 4.64 | -2.96 ** |
| Weight | 68.00 | 65.84 | 10.34 |  | 67.00 | 78.06 | 8.70 | -7.42 *** |
| Height | 69.00 | 171.90 | 6.98 |  | 68.00 | 182.51 | 6.44 | -9.24 *** |
| Alcohol drinks | 68.00 | 0.63 | 0.55 |  | 68.00 | 1.08 | 0.55 | -3.13 ** |
| How much commitment (0-9) | 64.00 | 5.92 | 1.64 |  | 65.00 | 5.72 | 1.43 | 0.73 |
| Time single in months | 62.00 | 38.69 | 65.31 |  | 59.00 | 37.03 | 65.45 | 0.14 |
|  | Average per cent | | | | | | | |
| Education | Female (N = 69) | | |  | Male (N = 69) | | |  |
| VMBO |  | 0.02 |  |  |  | 0.00 |  |  |
| HAVO |  | 0.04 |  |  |  | 0.02 |  |  |
| VWO |  | 0.02 |  |  |  | 0.15 |  |  |
| MBO |  | 0.13 |  |  |  | 0.08 |  |  |
| HBO |  | 0.30 |  |  |  | 0.38 |  |  |
| WO |  | 0.47 |  |  |  | 0.38 |  |  |

Note: *** *p* < .001, VMBO: the lowest completed high-school level, WO: the highest level (scientific education, Bachelor or Master degree)

**Table S10.** Comparisons (t-test) variables by sex

|  | Female | | |  | | Male | | | |
| --- | --- | --- | --- | --- | --- | --- | --- | --- | --- |
| Variable | M | SD |  | | M | | SD | t | P-Value |
| SDI | 4.25 | 1.47 |  | | 4.68 | | .93 | -1.692 | .094 |
| LSAS Fear | .46 | .49 |  | | .45 | | .64 | .099 | .921 |
| LSAS avoidance | .77 | .67 |  | | .81 | | .69 | -.271 | .787 |
| PANAS negative | 1.69 | .45 |  | | 1.72 | | .40 | -.280 | .780 |
| PANAS positive | 3.20 | .56 |  | | 3.41 | | .53 | -1.853 | .067 |
| Trust baseline | 7.10 | 1.31 |  | | 7.13 | | 1.10 | -.075 | .940 |
| Trust overall | 7.15 | 1.18 |  | | 7.11 | | .91 | .186 | .150 |
| *Note: There were no significant values found between male and females*  **Table S10** compares gender difference in Liebowitz Social Anxiety Scale (LSAS), Positive and Negative Affect Schedule (PANAS) and Sexual Desire Inventory (SDI). The SDI is comprised of 11 items about various sexual behaviours, on a 5-point Likert scale. The total score on the SDI is the sum of all 11 items, with higher scores reflecting a higher sexual desire. The LSAS is comprised of two subscales: performance and social interaction. The 24 questions ultimately lead to six subscale scores: total fear, fear of social interaction, fear of performance, total avoidance, avoidance of social interaction and avoidance of performance. The statements had to be answered on a 4-point scale (0 = not at all, 4 = totally).  The PANAS: consists of two 10-item mood scales, measuring positive affect (PA) and negative affect (NA). Participants are asked to rate their experience with a certain emotion on a 5-point scale (1 = very slightly or not at all, 5 = very much). | | | | | | | | | |

**Movie S1 (separate file). An example of physiological synchrony.** The video shows a nonverbal interaction where participants were instructed not to talk. At 00:04:00, the female will smile and the male partner reciprocates with a smile back. During this moment, we observe an increase in female’s and males’ skin conductance and heart rate (top two rows). Again, at 00:18:24, the female laughs; in response the male smiles and we again observe synchrony in heart rate and skin conductance. Thus, the purpose of the video is to explain how synchrony can occur. Importantly, not all smiles and laughs were paired with physiological synchrony, but in the case of this couple, they did. Further examination of these empirical visualizations suggested that physiological synchrony is more closely linked to “genuine” emotional exchange such as contagious smiles or uncontrolled laughter, as opposed to overt expressions used during polite communication (grins or nods).
